# Derivation and validation of a VISITAG™-guided contact force ablation protocol for pulmonary vein isolation

**DOI:** 10.1101/232694

**Authors:** David R. Tomlinson

## Abstract

**Aims** Following radiofrequency (RF) pulmonary vein isolation (PVI), atrial fibrillation (AF) recurrence mediated by recovery of pulmonary vein (PV) conduction is common. I examined whether comparative VISITAG™ (Biosense Webster Inc.) data analysis at sites showing intra-procedural recovery of PV conduction versus acutely durable ablation could inform the derivation of a more effective VISITAG™-guided contact force (CF) PVI protocol.

**Methods and results** Retrospective analysis of VISITAG™ Module annotated ablation site data in 10 consecutive patients undergoing CF-guided PVI without active VISITAG™ guidance. Employing 2mm positional stability range and lenient CF filter (force-over-time 10%, minimum 2g), inter-ablation site distance >10-12mm, adjacent 0g-minimum CF and short RF duration (3-5s) were associated with intra-procedural recovery of PV conduction. A VISITAG™-guided CF PVI protocol was derived employing ≤6mm inter-ablation site distances, minimum target ablation site duration ≥9s / force time integral (FTI) 100gs and 100% 1g-minimum CF filter. Seventy-two consecutive VISITAG™-guided CF PVI procedures were then undertaken using this protocol, with PVI achieved in all utilising 23.8[8.4] minutes total RF (30W, 48°C, 17ml/min, continuous RF application). Following protocol completion, acute intra-procedural spontaneous / dormant recovery of PV conduction requiring touch-up RF occurred in 1.4% / 1.8% of PVs, respectively. At 14[5] months’ follow-up in all 34 patients with paroxysmal AF ≥6 months’ post-ablation, 30 (88%) were free from atrial arrhythmia, off class I/III anti-arrhythmic medication.

**Conclusion** VISITAG™ provides means to identify and then avoid factors associated with intra-procedural recovery of PV conduction. This VISITAG™ Module-guided CF PVI protocol demonstrated excellent intra-procedural and long-term efficacy.

**Condensed abstract** Following CF-guided PVI, retrospective VISITAG™ Module analyses permitted the identification of ablation parameters associated with intra-procedural recovery of PV conduction. The derived VISITAG™ Module-guided CF PVI protocol employed short overall RF duration yet proved efficient at achieving PVI acutely, with long-term follow-up demonstrating high clinical efficacy.

## What’s New?

- The VISITAG™ Module (Biosense Webster Inc.) is the first tool providing automated and objective annotation of radiofrequency (RF) ablation data onto a 3D electroanatomical map.
- Ablation parameters associated with intra-procedural recovery of pulmonary vein (PV) conduction following contact force (CF)-guided PV isolation (PVI) were retrospectively identified, employing lenient VISITAG™ filters providing maximal RF data capture.
- A VISITAG™-guided CF PVI protocol was devised to eliminate adverse ablation parameters: Force-over-time 100﹪, minimum 1g, to eliminate intermittent catheter-tissue contact during RF; target ablation site duration (≥9s) / force time integral (≥100gs); ≤6mm inter-ablation site distance.
- This VISITAG™-guided CF PVI protocol proved highly effective both acutely and at long-term follow-up, yet with shorter RF duration than presently considered appropriate.
- The VISITAG™ Module represents an important advance in RF ablation guidance technology, permitting identification of adverse ablation features and their avoidance via objective annotation methodology.

## Introduction

Pulmonary vein isolation (PVI) represents the cornerstone of invasive treatment for symptomatic atrial fibrillation (AF).^1^ Improved procedural outcomes have resulted from the use of contact force (CF)-sensing radiofrequency (RF) ablation catheters, particularly when lesions are delivered with optimal CF.^2^ However, achieving durable PVI remains technically challenging, since late recovery of pulmonary vein (PV) conduction is commonly noted following index PVI even using current CF guidelines.^3^

The VISITAG™ Module (Biosense Webster Inc., Diamond Bar, CA) is the first technology permitting automated and objective ablation site annotation through catheter stability “filters” incorporating CF and positional data during RF application (figure 1). The ablation catheter position is measured every 16/17ms, from which is calculated the standard deviation (SD). Consecutive ablation site locations within a user-defined maximum range for the SD (in mm) are used to annotate a site meeting positional stability filter requirements. CF is measured every 50ms via the NaviStar^®^ THERMOCOOL^®^ SMARTTOUCH™ (ST) catheter (Biosense Webster Inc.). The CF filter applies a user-defined minimum CF for a percentage of RF time at sites meeting positional stability filter parameters. Consecutive catheter locations satisfying both positional and CF stability filter requirements for a minimum user-defined duration, result in automated ablation site annotation; a 3D and/or surface-projected tag indicates the mean catheter position at each site. RF duration is shown in an active count window (figure 1A), with ablation data allocation continuing until either the position or CF parameters exceed filter thresholds. Tag colouration can be used to indicate when user-defined targets have been achieved (e.g. displaying RF time, FTI or impedance drop). An icon indicates when VISITAG™ Module catheter stability filter thresholds are met, potentially aiding the attainment of user-defined parameters during RF (figures 1A-C). Respiration adjustment (ACCURESP™) may be applied, resulting in ablation data summation to the mean end-expiration site meeting filter criteria, although this has yet to be scientifically validated. Postablation, tags provide ablation site data including RF duration, mean and range of CF, FTI and impedance (figure 2). Presently, there are no guidelines on VISITAG™ Module use.

**Figure 1:**
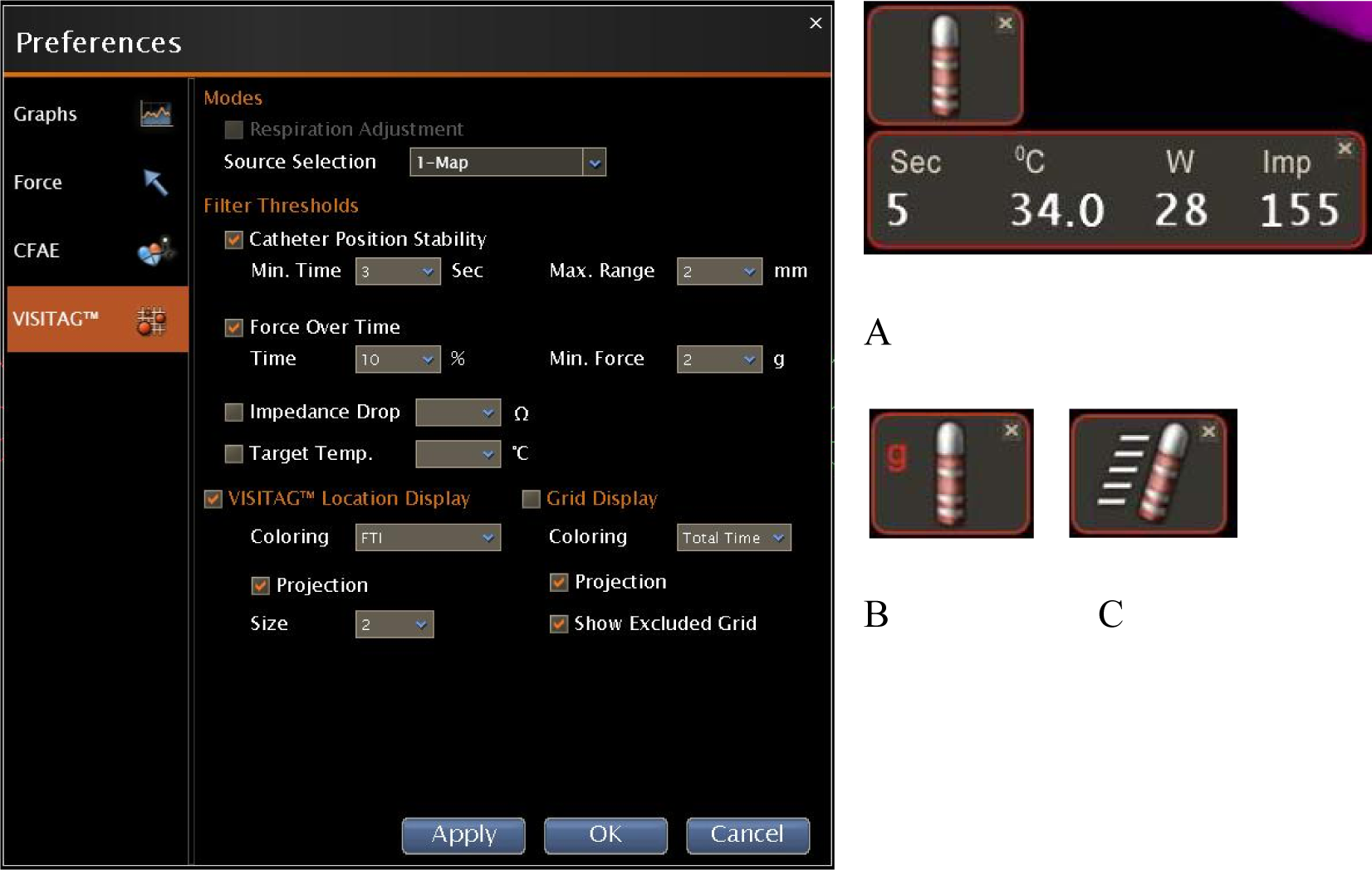
Derivation cohort VISITAG™ preferences. Catheter position stability and force-over-time filters are active, with parameters shown. VISITAG™ location display is “ON”; annotated tags set to 2mm radius, FTI colouration and surface projection “ON” (respiratory adjustment “OFF”). (A) Icon demonstrating that filter criteria are met during RF, with an active count of 5s; also shown are catheter temperature, RF power and impedance. In (B) and (C), icons demonstrate CF and positional stability out of range, respectively.

**Figure 2:**
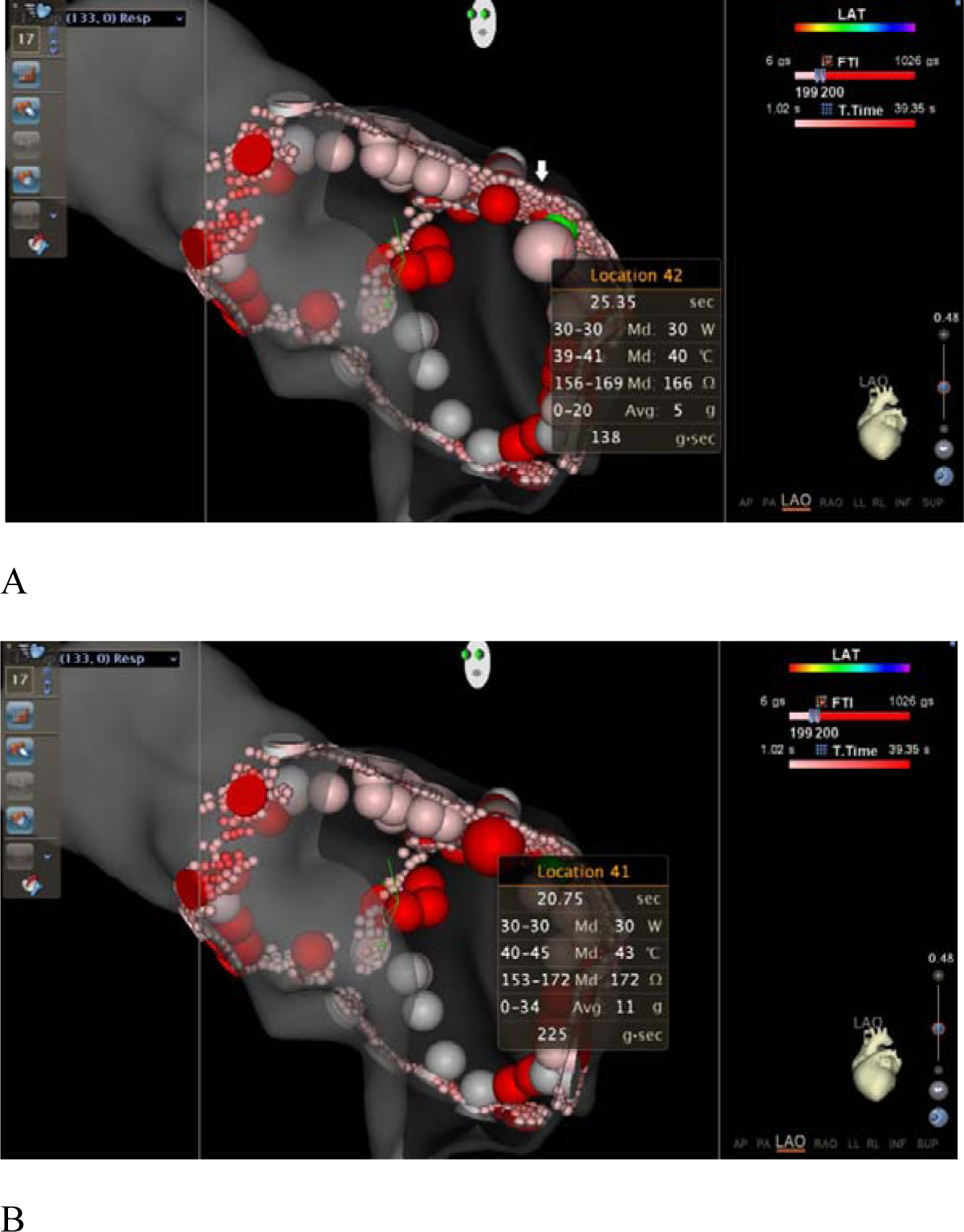
Clipping-plane of derivation cohort patient #7, right posterior carina site of spontaneous PV recovery. FTI colour transition at 200gs for illustrative purposes; 2mm tag radius. Adjacent 3D tags highlighted to display their parameters; on selection, the tag diameter increases. A green location tag marks the site of spontaneous recovery of PV conduction, with the red tag completing PVI seen in (A) under the arrow. The tag data view shows, from top to bottom: duration; range and median for each of power, temperature, impedance and CF; and total FTI.

This report describes VISITAG™ Module use initially as a recording tool for delivered RF ablation in a consecutive cohort of patients undergoing PVI. Retrospective analyses at sites of intra-procedural spontaneous and dormant recovery of PV conduction permitted the identification of adverse factors and derivation of a VISITAG™ Module-guided CF PVI ablation protocol. This was then employed in an unselected consecutive series of patients undergoing index PVI.

## Methods

The derivation cohort consisted of consecutive patients with symptomatic AF undergoing ablation according to treatment guidelines.^1^ Index CF-guided PVI was performed between July 2013 and January 2014, without prior personal use or knowledge of the VISITAG™ Module. Six procedures were performed before a 3-month period of ST catheter worldwide withdrawal. Interim ablation site data analysis was then performed, following which 4 further procedures were undertaken. Retrospective analyses for all 10 cases permitted derivation of a VISITAG™-guided CF PVI ablation protocol, employed during consecutive index PVI procedures from 30^th^ January 2014 to 11^th^ February 2016.

## Ablation protocol: Derivation cohort

General anaesthesia (GA) with endotracheal intubation and intermittent positive pressure ventilation was used in all. Double transseptal access was obtained with SL1 (St Jude Medical, Minneapolis, MN) and Agilis NxT sheaths (St Jude Medical). ACCURESP™ Module respiratory training was performed via a duodecapolar LASSO^®^ Nav (Biosense Webster Inc.) catheter in the left inferior PV and applied as required to the CARTO^®^3 geometry (version 3, Biosense Webster Inc.).

Electrograms were displayed on a BARD EP system (Boston Scientific Inc., Marlborough, MA,) at 200mm/s with 30 – 500Hz filter. Irrigated RF was performed in temperature-controlled mode (17ml/min, 48°C limit) at 30W with an F curve ST catheter via the Agilis sheath, ~1cm from the PV ostia, completing encirclement and PVI (entrance and exit block) using continuous RF. When considering an assessment of lesion completeness there was no strict adherence to bipolar electrogram attenuation or morphology changes, since RF artefact usually limited interpretation. Instead, RF applications were delivered in accordance with my “usual practice”, which targeted achieving RF duration of 10-20s at each site of ablation catheter stability, before moving to an immediately adjacent site. Target mean CF was 5-10g, except at sites where catheter stability was only achieved with lower CF. ST re-zeroing was performed following its withdrawal and re-introduction, or if an error was detected; i.e. mean CF ≥2g despite no endocardial tip contact. No technician ablation site annotation was used. Instead, objective VISITAG™ Module annotation was employed as 4mm diameter surface tags at sites meeting position and CF stability filter preferences. In order to maximise RF data capture for subsequent analysis and ensure that automated annotation would not indicate lesion completion (thereby eliminating bias), position stability duration was 3s (the minimum possible) and the force-over-time was 10% at 2g-minimum (i.e. permitting 90% RF at 0g). Arbitrarily 2mm positional stability range was used, since there were no pre-existing guidelines. FTI was not used to guide RF application, since FTI was not known to be an appropriate measure of ablation effect with VISITAG™. Neither the catheter stability icon nor the active tag count windows were used and ACCURESP™ respiratory adjustment was never applied to VISITAG™ filter preferences (figures 1A-C).

Following PVI, spontaneous recovery of PV conduction was assessed and eliminated during a 30-minute wait. Recovery sites were defined either by re-PVI (entrance and exit block) or alteration of LASSO^®^ activation occurring within 12s of antral RF opposite the LASSO^®^ electrode displaying earliest PV activation. Dormant recovery was evaluated and targeted 30 minutes after the last RF; during sinus rhythm, intravenous adenosine resulting in transient complete AV block was administered. If dormant recovery persisted, recovery sites were defined as above. Transient dormant recovery sites were defined by the elimination of this response following 1/2 adjacent <20s duration antral lesions delivered opposite the earliest LASSO^®^ PV electrogram noted. Outcomes were assessed following a 90-day blanking period, during which antiarrhythmic drug use (except amiodarone) was permitted. Clinic review with 12-lead ECG was at 3-4, 6-8, and 12-15 months. Freedom from atrial arrhythmia lasting >30s was assessed with symptom-driven ECG / ambulatory monitoring and 48 hour to 5 day ambulatory monitoring at 6-9 and 12-18 month intervals, or from pacemaker/ICD data when available.

## Ablation site data analysis

Using a proprietary export function, exported VISITAG™ data analyses were performed.Sites of spontaneous and/or dormant recovery of PV conduction sites were identified and comparisons made with cases showing acutely durable ablation at these same locations. Inter-ablation site distances were estimated visually with 3D tags displayed at 4mm diameter and represent the distance between centre-points of adjacent tags. Where tag overlap was ≥50%, data was summated, as a single site (at this time no precise distance measurement tool was available).

Statistical analyses were performed using GraphPad Prism version 4.03. Parametric data are expressed as mean [SD]; non-parametric data are presented as median (1^st^ – 3^rd^ quartile).Unpaired t test or the Mann Whitney test was used to assess statistical significance for continuous data, as appropriate. The modified Wald method was used to determine the confidence interval of a proportion. P <0.05 indicated a statistically significant difference. This work received Institutional Review Board (IRB) approval for publication as a description of an evaluation of clinical experience with a new ablation guidance product without guidelines on its use. All patients provided written, informed consent.

## Results

### Derivation cohort

Table 1 shows patient demographics, procedure and clinical outcome data. There were no complications and all PVs were isolated with a total RF duration of 22.3[5.8] minutes. Spontaneous and dormant recovery of PV conduction occurred in 8% of PVs; i.e. 6 sites.

**Table 1:**
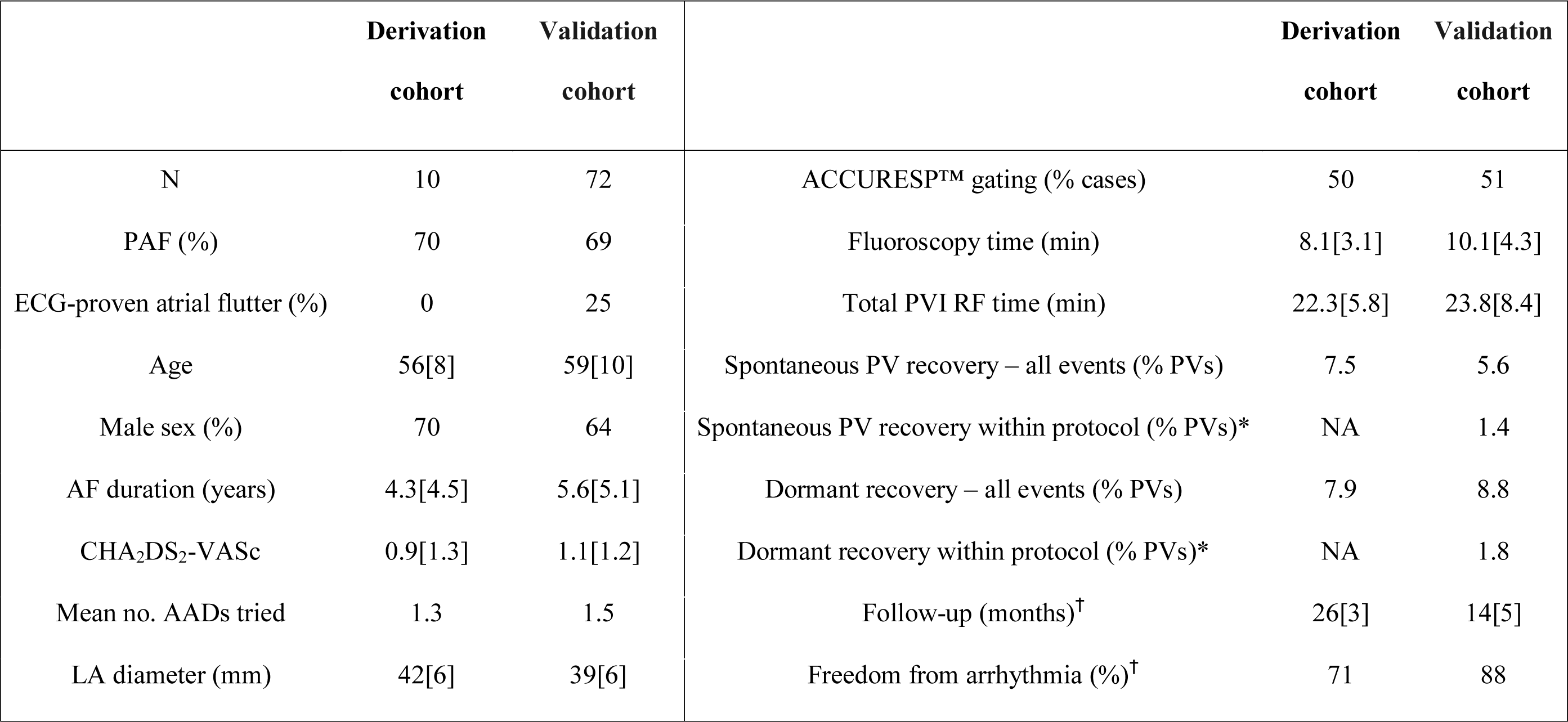
Patient demographics (left panel) and procedure and clinical outcome data (right panel). *Adjusted rate excludes recovery of PV conduction events due to unrecognised gaps / lower than target ablation delivery, or ST catheter zeroing errors; ^†^data for all PAF patients with >6 months’ follow-up data (AADs, antiarrhythmic drugs; PAF, paroxysmal AF).

### Ablation site data: Derivation cohort

595 annotated ablation sites were analysed, isolating 20 PVs (i.e. 5 of the first 6 cases; data file corruption prevented complete analysis for patient #4). Ablation sites were closely-spaced – 92% were within 4mm of their nearest tag. Left PVI ablation sites had shorter duration (4.9s versus 6.3s, p=0.007), lower mean CF (6.4g versus 11.5g, p<0.0001) and lower FTI (40gs versus 80gs, p<0.0001) than right-sided PVI sites (table 2). Intermittent catheter contact was more common during left PV ablation – 31% compared to 12% (p<0.0001). Using the TOCCATA 7 segment PV model,^4^ the median total FTI per segment was 432gs and 838gs for the left and right PV pair, respectively (p=0.02).

**Table 2:**
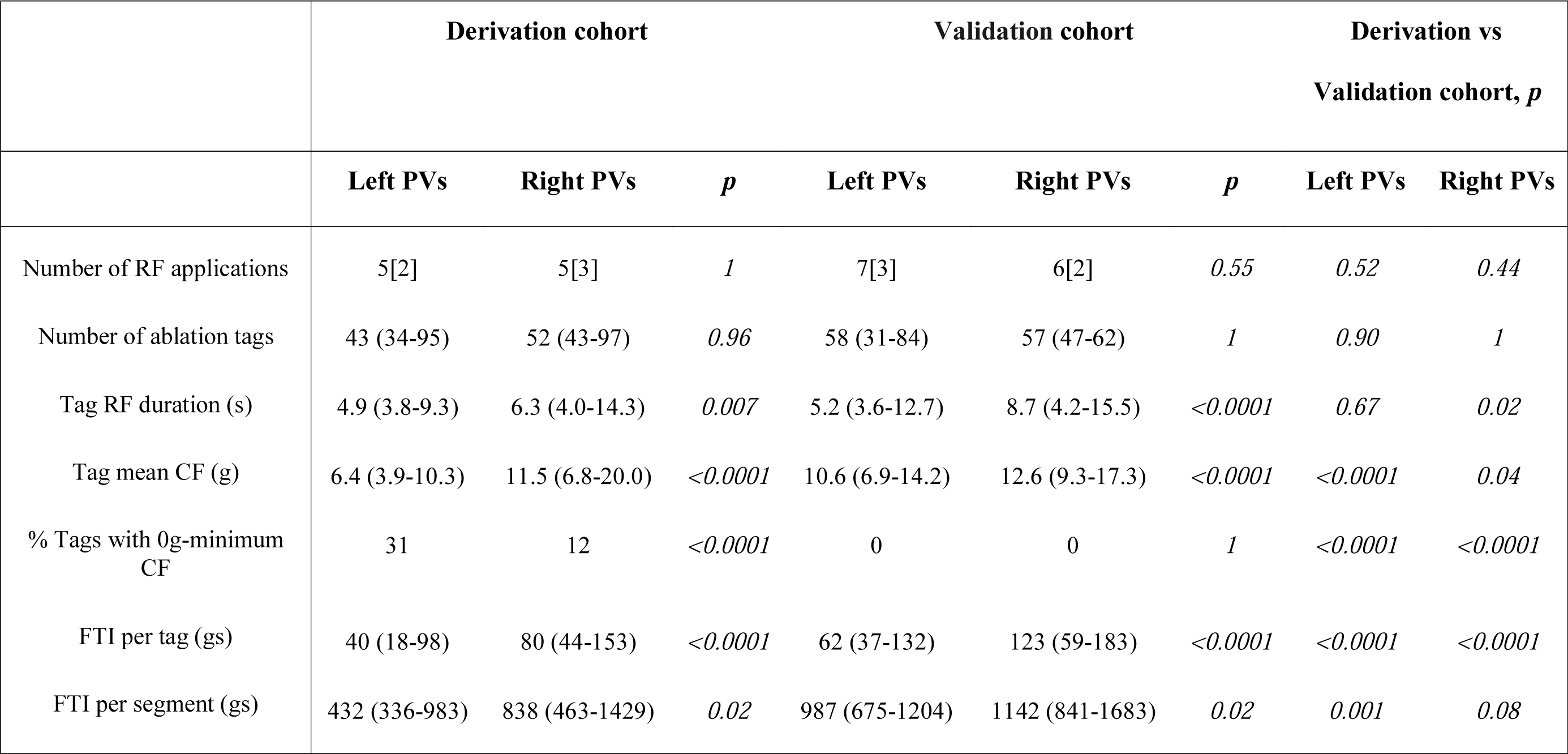
Exported VISITAG™ data, with comparative analyses within and between cohorts according to each vein pair.

|  | Derivation cohort |  |  | Validation cohort |  |  | Derivation vs<br>Validation cohort, *p* |  |
| --- | --- | --- | --- | --- | --- | --- | --- | --- |
|  | Left PVs | Right PVs | *p* | Left PVs | Right PVs | *p* | Left PVs | Right PVs |
| Number of RF applications | 5[2] | 5[3] | *1* | 7[3] | 6[2] | *0.55* | *0.52* | *0.44* |
| Number of ablation tags | 43 (34-95) | 52 (43-97) | *0.96* | 58 (31-84) | 57 (47-62) | *1* | *0.90* | *1* |
| Tag RF duration (s) | 4.9 (3.8-9.3) | 6.3 (4.0-14.3) | *0.007* | 5.2 (3.6-12.7) | 8.7 (4.2-15.5) | *<0.0001* | *0.67* | *0.02* |
| Tag mean CF (g) | 6.4 (3.9-10.3) | 11.5 (6.8-20.0) | *<0.0001* | 10.6 (6.9-14.2) | 12.6 (9.3-17.3) | *<0.0001* | *<0.0001* | *0.04* |
| % Tags with 0g-minimum CF | 31 | 12 | *<0.0001* | 0 | 0 | *1* | *<0.0001* | *<0.0001* |
| FTI per tag (gs) | 40 (18-98) | 80 (44-153) | *<0.0001* | 62 (37-132) | 123 (59-183) | *<0.0001* | *<0.0001* | *<0.0001* |
| FTI per segment (gs) | 432 (336-983) | 838 (463-1429) | *0.02* | 987 (675-1204) | 1142 (841-1683) | *0.02* | *0.001* | *0.08* |

Inter-ablation site distance >10-12mm on the left atrial appendage ridge, adjacent sites below the right inferior PV carina with 0g-minimum CF and adjacent short duration sites (3-5s) with 3-7mm inter-ablation site distances at 4 right PV locations were associated with intra-procedural recovery of PV conduction (table 3 and e.g. figure 2). In comparison, acutely durable ablation effect in the other 9 cases was always seen in association with inter-ablation site distance ≤6mm, no adjacent 0g minimum CF sites and RF duration >6s.

**Table 3:** Intra-procedural recovery of PV conduction site characteristics for derivation cohort (LAA, left atrial appendage; RSPV, right superior PV; RIPV, right inferior PV; T, tag duration; Min, minimum). Note: Carina recovery of conduction sites were all within 10mm of the carina.

| Patient | PV site | Recovery type | Inter-ablation site<br>distance (mm) | Immediately adjacent tag data |  |  |  |  |  |
| --- | --- | --- | --- | --- | --- | --- | --- | --- | --- |
|  |  |  |  | T (s) | FTI (gs) | Min CF (g) | T (s) | FTI (gs) | Min CF (g) |
| 4 | LAA ridge | Dormant | 10-12 | 5.4 | 23 | 1 | 24.0 | 131 | 1 |
| 7 | Posterior RIPV below carina | Spontaneous | 7 | 25.4 | 138 | 0 | 20.8 | 225 | 0 |
| 10 | RSPV roof | Spontaneous | 6 | 3.0 | 24 | 3 | 5.1 | 76 | 6 |
| 10 | Posterior RIPV below carina | Spontaneous | 7 | 3.0 | 26 | 1 | 3.0 | 64 | 1 |
| 10 | Anterior RSPV | Dormant | 3 | 3.0 | 48 | 4 | 3.0 | 50 | 4 |
| 10 | Posterior RSPV above carina | Dormant | 3 | 3.0 | 48 | 4 | 3.0 | 50 | 4 |

### Follow-up

At 26[3] months’ follow-up in all 7 patients with paroxysmal AF (PAF), 5 were atrial arrhythmia-free, off class I/III anti-arrhythmic medication.

### VISITAG™-guided CF PVI: Protocol derivation

To eliminate adjacent 0g-minimum CF ablation sites (i.e. recovery of PV conduction event for patient #7, table 3), a force-over-time filter of 100% 1g-minimum was applied, so ablation site annotation would only occur during constant catheter-tissue contact. Since 92% of ablation sites were within 4mm of their closest site and as the median RF duration for left PV ablation sites was 4.9s, I considered a minimum RF duration ≥9s over a ≤6mm inter-ablation site distance to be appropriate during continuous RF application (i.e. recovery of PV conduction events for patients #4 and #10, table 3). For “RF ON” sites, 4-5 s additional RF was utilised to compensate for power increase from 0W. For anterior PV wall sites without risk of oesophageal thermal trauma, the minimum target ablation site parameters included FTI ≥100gs when this resulted in greater ablation site duration. When carina ablation was required to achieve PVI, the target minimum ablation site duration was 15-20s / FTI 200gs – which ever resulted in greatest duration – in view of known greater carina wall thickness.^5^ The minimum display duration remained 3s, for rapid annotation and maximal ablation data display; positional stability range remained 2mm.

During protocol validation, technical differences from the derivation cohort procedures were limited to VISITAG™ Module-CARTO^®^3 interface use: (1) Catheter stability icon was used to optimise stability attainment during RF; (2) tag colour change (FTI) and the tag count window (RF duration) were used to determine attainment of target parameters (figures 1A and 3); (3) careful CF waveform inspection helped avoid 0g-minimum CF; (4) inter-ablation site distances and RF parameters were assessed using 10% CARTO^®^3 map transparency following PVI, with additional RF closing any visible gaps between 3D tags, as required; (5) consecutive tags with ≥50% overlap were considered to represent ablation at a single site, with RF data summated.

**Figure 3:**
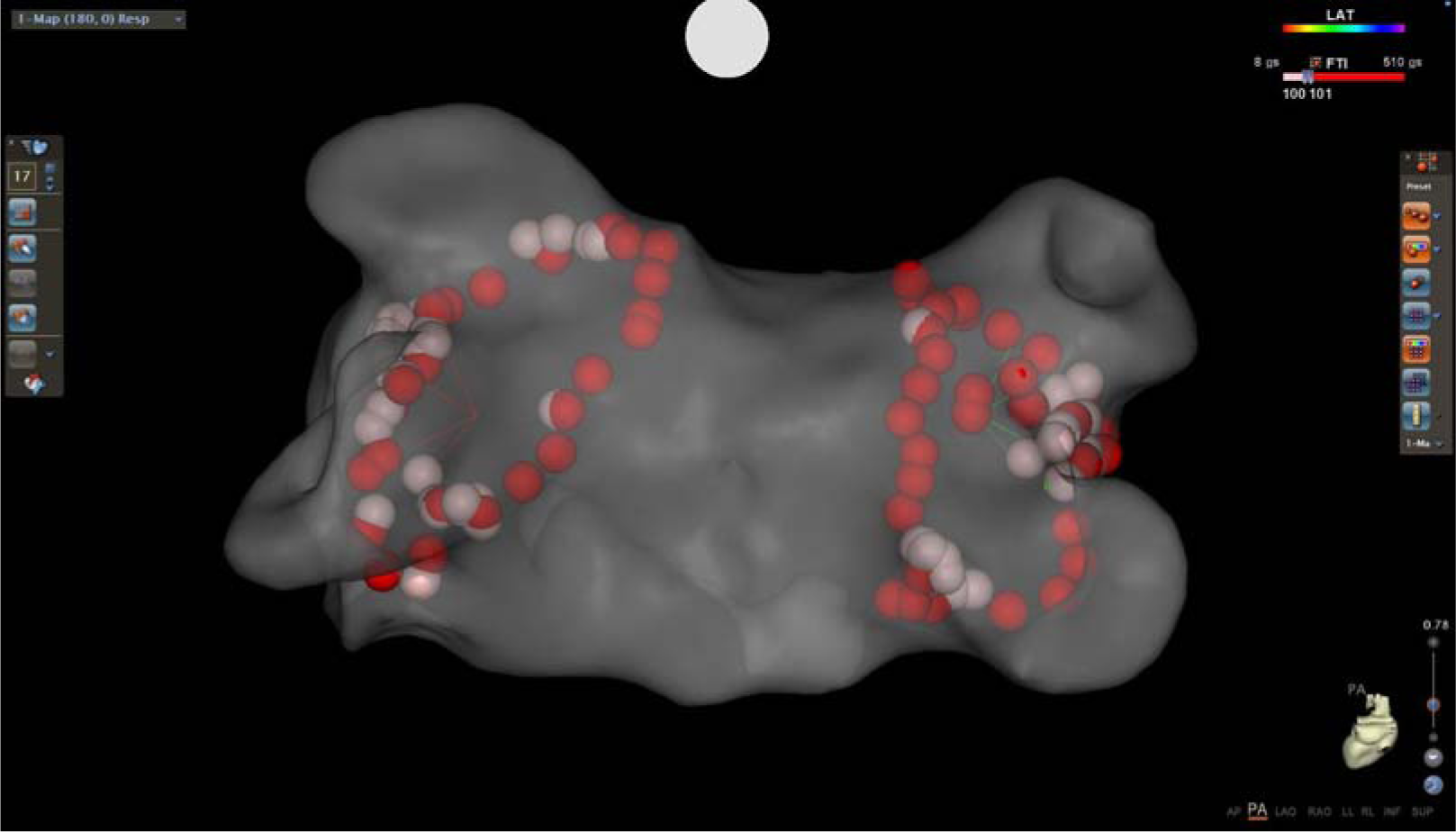
PA view of index PVI annotated lesion set for patient #29, in whom enduring PVI was found at redo. Tags are set to 4mm diameter, with transition to red at 101gs FTI (map transparency, 10%). The maximum FTI of 510gs resulted from 19.3s RF at 26.5g mean CF at the right PV carina.

### Validation cohort: Outcomes

In 72 consecutive patients, PVI was achieved in 100% (288 veins) without complication; table 1 shows demographic, procedure and clinical outcome data. The total RF duration was 23.8[8.4] minutes; spontaneous and dormant recovery of PV conduction occurred in 5.6% and 8.8% of PVs, respectively; i.e. 16 spontaneous and 25 dormant recovery events. Of these, 8 spontaneous and 16 dormant PV recovery events were due to unidentified errors in deployment of the VISITAG™-guided CF PVI protocol: Unrecognised >6mm inter-ablation site gaps accounted for all 8 spontaneous and 10 dormant PV recovery events; unrecognised short duration RF resulted in 6 dormant PV recovery events. ST catheter zeroing error noted intra-procedurally was associated with 4 spontaneous and 4 dormant PV recovery events in 6 patients. Therefore, only 4 spontaneous and 5 dormant recovery events occurred following completion of the intended VISITAG™-guided CF PVI protocol – i.e. 1.4% and 1.8% PVs, respectively. Of these, 3 spontaneous and 2 dormant recovery events occurred at the carina, where carina ablation had not yet been performed. Consequently, intentional RF delivery greater than that of the VISITAG™-guided CF PVI protocol was required to eliminate 1 spontaneous and 3 dormant PV recovery events – i.e. 0.3% and 1.1% PVs, respectively. One patient (#15) required RF delivery exceeding the VISITAG™-guided CF protocol in order to achieve initial PVI; 400 – 800gs FTI tags were required at multiple sites to the left superior and both right PVs. Uniquely, this patient also had myelofibrosis with splenomegaly; posterior left atrial wall thickness 2.5mm on cardiac MRI.

### Ablation site data: Validation cohort

1049 ablation site tags isolating 36 PVs were analysed; i.e. 9 of the first 13 PAF cases; the first 2 were excluded (possible learning curve) as well as patient #15 and a case with zeroing error. Left PVI sites had shorter duration (5.2s versus 8.7s, p<0.0001), lower mean CF (10.6g versus 12.6g, p<0.0001) and lower FTI (62gs versus 123gs, p<0.0001) than right PV sites (table 2). The median total FTI per segment was 987gs and 1142gs for the left and right vein pairs, respectively (p=0.02). Compared to the derivation cohort, VISITAG™-guided CF PVI ablation sites had greater mean CF and for the right PV, and RF duration was also significantly increased. Consequently, there was a significantly greater ablation site FTI for both PVs – from 40gs to 62gs for the left PV and from 80gs to 123gs for right PV (p<0.0001).

### Follow-up

At 14[5] months’ follow-up in all 34 paroxysmal AF patients with ≥6 months’ follow-up, 30 (88%) were free from atrial arrhythmia, off class I/III anti-arrhythmic medication. Of the PVI failures: Patient #2 required flecainide (50mg bd) for ventricular premature beat suppression, but was AF-free over 25 months’ follow-up (dual chamber ICD data); #19 had 6% PAF burden on pacemaker interrogation at 13 months’ follow-up, off antiarrhythmic medication, with redo not required due to symptom improvement (pre-ablation, 46% AF); #25 had 1 episode of typical atrial flutter at 6 months’ post-ablation (index PVI and flutter ablation), but no AF at 13 months; #29 had post-ablation PAF and underwent redo 7 months later but with all PVs demonstrating durable isolation without dormant recovery (figure 3). For this case the annotated index PVI median ablation site FTI, duration and mean CF were 103gs (50-184gs), 6.6s (3.7-14.9s) and 13.0g (8.2-16.4g), respectively.

## Discussion

This is the first reported use of the VISITAG™ Module as a retrospective “investigational tool” following CF-guided PVI, demonstrating how factors resulting in spontaneous and/or dormant recovery of PV conduction may be determined. In a consecutive series of patients undergoing PVI, lenient VISITAG™ stability filter parameters were used for automated annotation, maximising RF ablation data capture and eliminating bias. Retrospective analyses identified spontaneous and/or dormant intra-procedural recovery of PV conduction events where adjacent ablation sites either exceeded a critical distance (>10-12mm), showed intermittent catheter-tissue contact (i.e. 0g-minimum CF) or short RF duration (3-5s). This report also demonstrates how the VISITAG™ Module may be used to eliminate these parameters, through: (1) A force-over-time filter eliminating intermittent catheter-tissue contact during ablation site annotation (100%, 1g-minimum); (2) an icon alert for suboptimal catheter stability parameters; (3) annotated site colouration and RF count window identifying minimum target parameters; and (4) a means to assess inter-ablation site distances. The resulting VISITAG™-guided CF PVI protocol proved highly effective, with intra-procedural recovery of PV conduction in <2% of PVs and mid-term clinical success in 88% patients with paroxysmal AF.

These outcomes good outcomes resulted from overall short RF duration and both lower CF and FTI than presently considered appropriate. Indeed, the FTI delivered per segment in this present study was equivalent to a single ablation site in TOCCATA.^4^ Based upon these findings, the current guidelines of 20g target CF, with absolute minimum of 10g and absolute minimum 400gs FTI per ablation lesion during PVI^6^ should be viewed with caution. Clearly, without objective means of ablation site annotation according to pre-defined levels of catheter stability, any attribution of treatment effect to an “ablation site” is prone to error. In TOCCATA the definition of “valid” CF records for analysis were those “with stable CF over 15 consecutive seconds”.^4^ However, CF “stability” (however that is defined) has never been proven to equate with positional stability *in vivo.* Therefore, compared to data derived from VISITAG™-based objective annotation, important methodological flaws exist for all current published guidelines regarding the optimal CF/FTI or impedance change required to achieve a completed ablation effect *in vivo*.^3,4,6–12^

One publication to-date describes acute outcomes of VISITAG™ guidance during PVI. In 42 patients under GA (jet ventilation +/− periods of apnoea), point-by-point PVI was performed using a non-CF sensing catheter via an Agilis sheath. VISITAG™ was used to annotate sites at >15s duration, when an impedance decrease of ≥5% (7-8Ω) occurred over this time, within a 2mm (SD) range. Compared to the previous 42 PVI procedures with subjective annotation, VISITAG™ use resulted in a significant decrease in intra-procedural recovery of PV conduction (6% versus 25% vein pairs). However, there was no difference in the RF times between these groups – 49.9 and 50.2 minutes for VISITAG™ and operator annotation, respectively.^13^

Evidence in support of a short RF duration approach to PVI comes from a number of sources. Firstly, the atrial wall is thin. Cadaveric studies demonstrated thickest PV myocardial sleeves at the carina (1.6-2.0mm), with myocardial thickness <1.3mm at all other veno-atrial sites.^5^ Furthermore, computed tomography (CT) assessment of left atrial muscular wall thickness prior to ablation of persistent AF demonstrated a mean thickness of 1.89mm (range 0.5-3.5mm).^14^ Secondly, preparations of irrigated RF using “conventional” RF duration, result in lesions deeper than required *in vivo*. For example, in a bovine skeletal muscle model of intermittent catheter contact at 20W, an FTI of 200gs (CF range 0-22g) resulted in lesions of 4mm depth.^15^ Finally, unipolar electrogram (UE) markers of histologically confirmed transmural atrial lesions occur rapidly after the onset of irrigated RF. Following *in vivo* canine right atrial ablation using power-controlled RF at 30W (17ml/min irrigation, 48°C) with a ST catheter, RF durations according to unipolar R wave completion – a histologically validated marker of transmural (TM) atrial lesions^16^ – were compared with conventional 30s duration applications. All lesions were delivered with a mean CF of 11g. In 135 lesions with RF termination at R wave completion (i.e. R+0s), 95% were TM (mean depth 4.3mm). In 28 lesions of 30s duration, 100% were transmural (mean depth 5.9mm). However, while extracardiac thermal trauma was never seen at R+0s lesions, 10.7% of 30s duration sites demonstrated extra-cardiac thermal trauma. Notably, the mean time to R wave completion in this study was 7s.^17^

### Study limitations

Invasive assessment is required to prove that VISITAG™-guided CF PVI represents an optimal means to achieve durable PVI. Furthermore, trans-telephonic monitoring was not used, reflecting this report’s nature as an ablation guidance product evaluation during my usual clinical practice. This VISITAG™-guided CF PVI ablation protocol is only applicable to a single operator, using the techniques described. However, in view of the detailed identification of delivered ablation lesion sets now possible through VISITAG™, any operator working within the CF and positional stability ranges described and using continuous RF application at 30W, may be expected to replicate these ablation site data and clinical outcomes. Alternatively, and scientifically more appropriately, any operator can identify parameters differentiating between acutely durable PVI and sites of recovery of PV conduction using their usual techniques in order to derive “personalised” VISITAG™-guided CF PVI protocols of theoretically greater efficacy. These hypotheses clearly require further study for confirmation.

The utility of VISITAG™ respiratory adjustment is not known. However, respiratory adjustment requires 2 respiratory cycles for automated ablation site annotation, resulting in 7-8s delay to automated annotation even when using 3s stability duration. This may hinder the attainment of target short RF duration ablation site parameters through a lack of early feedback to the operator of sites meeting catheter stability requirements. Insufficient ST catheter zeroing errors occurred to identify causative factors, although since May 2015 the ST catheter now has auto-error detection and correction (Noam Seker-Gafni, personal communication). However, unnoticed zeroing errors may have occurred during this study, so the reported mean CF and FTIs may be greater than those actually delivered.

During VISITAG™-guided CF PVI, some inadvertently overlapping 3D-located ablation sites occurred, resulting in greater than target focal RF delivery and possibly contributing to procedural outcomes (e.g. figure 3). Furthermore, visual assessment of inter-ablation site distances is prone to possible error, resulting in greater than target inter-ablation site gaps. However, there was no systematic attempt to achieve target inter-ablation site distances differing from those stated in the derived VISITAG™-guided CF PVI protocol. Finally, this report does not prove that FTI represents a suitable ablation parameter to guide RF delivery for all operators, since 100gs FTI may simply represent a “personal sweet spot” for ≥9s ablation site duration, when operating at 11-13g mean CF and constant catheter-tissue contact.

### Conclusions

The VISITAG™ Module represents an important advance in ablation guidance technology, providing both a means to identify and avoid factors associated with intra-procedural recovery of PV conduction. A derived VISITAG™-guided CF PVI protocol employed short RF duration and lower FTI than presently considered appropriate, yet was highly effective at achieving PVI, excellent clinical outcomes.

## Acknowledgments

I am grateful to Ian Lines and Cherith Wood, Cardiac Physiologists, for their technical support into all cases conducted in this report. I am also grateful to Jon Toms (Biosense Webster) for his technical assistance with the initial VISITAG™ cases, and Tal Bar-on, Noam Seker-Gafni, Einav Geffen, Assaf Rubissa and colleagues at the Haifa Technology Center, Israel for their assistance with VISITAG™ Module exported data extraction for the evaluation cohort and their review of my technical description of VISITAG™ Module.

## Conflict of interest

none declared.

## References

1. Calkins H, Kuck KH, Cappato R, Brugada J, Camm AJ, Chen S-A, et al. 2012 HRS/EHRA/ECAS expert consensus statement on catheter and surgical ablation of atrial fibrillation: recommendations for patient selection, procedural techniques, patient management and follow-up, definitions, endpoints, and research trial design. Heart Rhythm [Internet]. 2012 Apr [cited 2015 Aug 15];9(4):632–96.e21.

2. Reddy VY, Dukkipati SR, Neuzil P, Natale A, Albenque J-P, Kautzner J, et al. Randomized, Controlled Trial of the Safety and Effectiveness of a Contact Force-Sensing Irrigated Catheter for Ablation of Paroxysmal Atrial Fibrillation: Results of the TactiCath Contact Force Ablation Catheter Study for Atrial Fibrillation (TOCCASTAR) S. Circulation [Internet]. 2015 Sep 8 [cited 2015 Sep 30];132(10):907–15.

3. Kautzner J, Neuzil P, Lambert H, Peichl P, Petru J, Cihak R, et al. EFFICAS II: optimization of catheter contact force improves outcome of pulmonary vein isolation for paroxysmal atrial fibrillation. Europace [Internet]. 2015 Aug [cited 2015 Nov 11];17(8):1229–35.

4. Reddy VY, Shah D, Kautzner J, Schmidt B, Saoudi N, Herrera C, et al. The relationship between contact force and clinical outcome during radiofrequency catheter ablation of atrial fibrillation in the TOCCATA study. Heart Rhythm [Internet]. 2012 Nov [cited 2015 Oct 19];9(11):1789–95.

5. SY Ho, JA Cabrera, VH Tran, J Farre, RH Anderson DS-Q. Architecture of the pulmonary veins: relevance to radiofrequency ablation. Heart. 2001;86:265–70.

6. Neuzil P, Reddy VY, Kautzner J, Petru J, Wichterle D, Shah D, et al. Electrical reconnection after pulmonary vein isolation is contingent on contact force during initial treatment: results from the EFFICAS I study. Circ Arrhythm Electrophysiol [Internet]. 2013 Apr [cited 2015 Oct 19];6(2):327–33.

7. Reichlin T, Knecht S, Lane C, Kühne M, Nof E, Chopra N, et al. Initial impedance decrease as an indicator of good catheter contact: insights from radiofrequency ablation with force sensing catheters. Heart Rhythm [Internet]. 2014 Feb [cited 2015 Nov 11];11 (2): 194–201.

8. Kumar S, Chan M, Lee J, Wong MCG, Yudi M, Morton JB, et al. Catheter-tissue contact force determines atrial electrogram characteristics before and lesion efficacy after antral pulmonary vein isolation in humans. J Cardiovasc Electrophysiol [Internet]. 2014 Feb [cited 2015 Oct 19];25(2):122–9.

9. Ullah W, Hunter RJ, Baker V, Dhinoja MB, Sporton S, Earley MJ, et al. Target indices for clinical ablation in atrial fibrillation: insights from contact force, electrogram, and biophysical parameter analysis. Circ Arrhythm Electrophysiol [Internet]. 2014 Feb [cited 2015 Nov 8];7(1):63–8.

10. Park C-I, Lehrmann H, Keyl C, Weber R, Schiebeling J, Allgeier J, et al. Mechanisms of Pulmonary Vein Reconnection After Radiofrequency Ablation of Atrial Fibrillation: The Deterministic Role of Contact Force and Interlesion Distance. J Cardiovasc Electrophysiol [Internet]. 2014 Jul 2 [cited 2015 Oct 19];25(7):701–8.

11. Ullah W, Hunter RJ, Baker V, Dhinoja MB, Sporton S, Earley MJ, et al. Factors affecting catheter contact in the human left atrium and their impact on ablation efficacy. J Cardiovasc Electrophysiol [Internet]. 2015 Mar [cited 2016 Jan 31];26(2):129–36.

12. Knecht S, Reichlin T, Pavlovic N, Schaer B, Osswald S, Sticherling C, et al. Contact force and impedance decrease during ablation depends on catheter location and orientation: insights from pulmonary vein isolation using a contact force-sensing catheter. J Interv Card Electrophysiol [Internet]. 2015 Sep [cited 2015 Nov 11];43(3):297–306.

13. Anter E, Tschabrunn CM, Contreras-Valdes FM, Buxton AE, Josephson ME. Radiofrequency ablation annotation algorithm reduces the incidence of linear gaps and reconnection after pulmonary vein isolation. Heart Rhythm [Internet]. 2014 May [cited 2015 Nov 11];11(5):783–90.

14. Beinart R, Abbara S, Blum A, Ferencik M, Heist K, Ruskin J, et al. Left atrial wall thickness variability measured by CT scans in patients undergoing pulmonary vein isolation. J Cardiovasc Electrophysiol. 2011;22(11):1232–6.

15. Shah DC, Lambert H, Nakagawa H, Langenkamp A, Aeby N, Leo G. Area under the real-time contact force curve (force-time integral) predicts radiofrequency lesion size in an in vitro contractile model. J Cardiovasc Electrophysiol [Internet]. 2010 Sep [cited 2015 Nov 11];21(9): 1038–43.

16. Otomo K, Uno K, Fujiwara H, Isobe M IY. Local unipolar and bipolar electrogram criteria for evaluating the transmurality of atrial ablation lesins at different catheter orientations relative to the endicardial surface. Heart Rhythm. 2010;7(9):1291–300.

17. Bortone A, Brault-Noble G, Appetiti A, Marijon E. Elimination of the negative component of the unipolar atrial electrogram as an in vivo marker of transmural lesion creation: acute study in canines. Circ Arrhythm Electrophysiol [Internet]. 2015 Aug [cited 2015 Nov 23];8(4):905–11.

